# Focal DNA hypo-methylation in cancer is mediated by transcription factors binding

**DOI:** 10.1101/2021.04.20.440687

**Authors:** Dylane Detilleux, Yannick G Spill, Delphine Balaramane, Michaël Weber, Anaïs Flore Bardet

**Affiliations:** CNRS, University of Strasbourg, UMR7242 Biotechnology and Cell Signaling, Illkirch 67412, France

**Author notes:** Shared first authors (Dylane Detilleux, Yannick G Spill).

## Abstract

Aberrant DNA methylation has emerged as a hallmark of cancer cells and profiling their epigenetic landscape has widely been carried out in many types of cancer. However, the mechanisms underlying changes in DNA methylation remain elusive. Transcription factors, initially thought to be repressed from binding by DNA methylation, have recently emerged as potential drivers of DNA methylation patterns. Here we perform a rigorous bioinformatic analysis integrating the massive amount of data available from The Cancer Genome Atlas to identify transcription factors driving aberrant DNA methylation. We predict TFs known to be involved in cancer as well as novel candidates to drive hypo-methylated regions such as FOXA1 and GATA3 in breast cancer, FOXA1 and TWIST1 in prostate cancer and NFE2L2 in lung cancer. We also predict TFs that lead to hyper-methylated regions upon TF loss such as EGR1 in several cancer types. Finally, we validate experimentally that FOXA1 and GATA3 mediate hypo-methylated regions in breast cancer cells. Our work shows the importance of TFs as upstream regulators shaping DNA methylation patterns in cancer.

## INTRODUCTION

DNA methylation is the most studied epigenetic modification in cancer^1^. Hyper-methylation of CpG island promoters of tumor suppressor genes associated to gene silencing is a hallmark of cancer^2^. More recently, regions of global hypo-methylation were identified in cancer as long partially methylated domains observed at late-replicating lamina-associated domains^3,4^.

Methylation of DNA occurs on cytosines mostly within CpG dinucleotides and is catalysed by DNA methyltransferases: DNMT1, DNMT3a and DNMT3b. DNA methylation is abundant throughout the genome except at CpG islands that are constitutively protected from DNA methylation. Initially, DNA methylation has been described as a transcriptional repressor, where the presence of DNA methylation at gene promoters would block transcription factor (TF) binding leading to gene silencing^5,6^. More recently, genome-wide studies showed that active regulatory elements bound by TFs invariably correlate with focal regions of low methylation and in contrast to the classical model, showed that TF binding could induce active demethylation mediated by the TET enzymes^7–10^. This strong anti-correlation between patterns of TF binding and DNA methylation therefore enables to infer active regulatory regions using DNA methylation data. However, it does not provide causal information about which of TF binding or DNA methylation regulates one another.

Despite extensive studies, the mechanisms leading to the accumulation of aberrant DNA methylation patterns in cancer are still poorly understood. Recent profiling of chromatin accessible regions marking TF occupancy in primary cancer samples correlated them to hypo-methylated regions suggested to be driven by key TFs^11^. However, investigating TF binding in primary cancer samples at a large scale remains challenging due to technical limitations.

Here, we exploited the massive amount of primary DNA methylation datasets from The Cancer Genome Atlas (TCGA) to predict TFs driving aberrant DNA methylation in cancer. We first identified differentially methylated regions (DMRs) between cancer and healthy samples. We performed a TF motif enrichment analysis to predict TF binding in DMRs. We then integrated matching TF expression data to distinguish TFs expected to drive DNA methylation changes in cancer. Finally, we validated our predictions in breast cancer cells and showed that FOXA1 and GATA3 indeed mediate DNA hypo-methylation.

## RESULTS

### Processing of the TCGA methylation data

To study DNA methylation changes in cancer, we retrieved 8425 raw methylation datasets, generated from Illumina Infinium HumanMethylation450 BeadChip (HM450), for 32 available cancer types from the TCGA resource. We processed the data with the ChAMP pipeline^12,13^ and performed normalization using noob^14^ as implemented in minfi^15,16^.

We trained a quadratic discriminant analysis aiming to classify the cancer and healthy samples in two groups for each cancer type and discarded samples that were misclassified (**Figure 1a**). Based on the number and dispersion of samples, we retained 13 cancer types for further analysis: Bladder urothelial carcinoma (BLCA), Breast invasive carcinoma (BRCA), Cholangiocarcinoma (CHOL), Colon adenocarcinoma (COAD), Head and neck squamous cell carcinoma (HNSC), Kidney renal clear cell carcinoma (KIRC), Kidney renal papillary cell carcinoma (KIRP), Liver hepatocellular carcinoma (LIHC), Lung adenocarcinoma (LUAD), Lung squamous cell carcinoma (LUSC), Prostate adenocarcinoma (PRAD), Rectum adenocarcinoma (READ), Thyroid carcinoma (THCA) and Uterine corpus endometrial carcinoma (UCEC). In the case of BRCA, representative of other cancer types (**Figure 1a**), we observe distinct clustering of the healthy and cancer samples and an expected broader heterogeneity of cancer samples. The results of this analysis can be visualized interactively for all cancer types on our webserver http://bardet.u-strasbg.fr/cancermethtf/ in the section “Data”.

**Figure 1:**
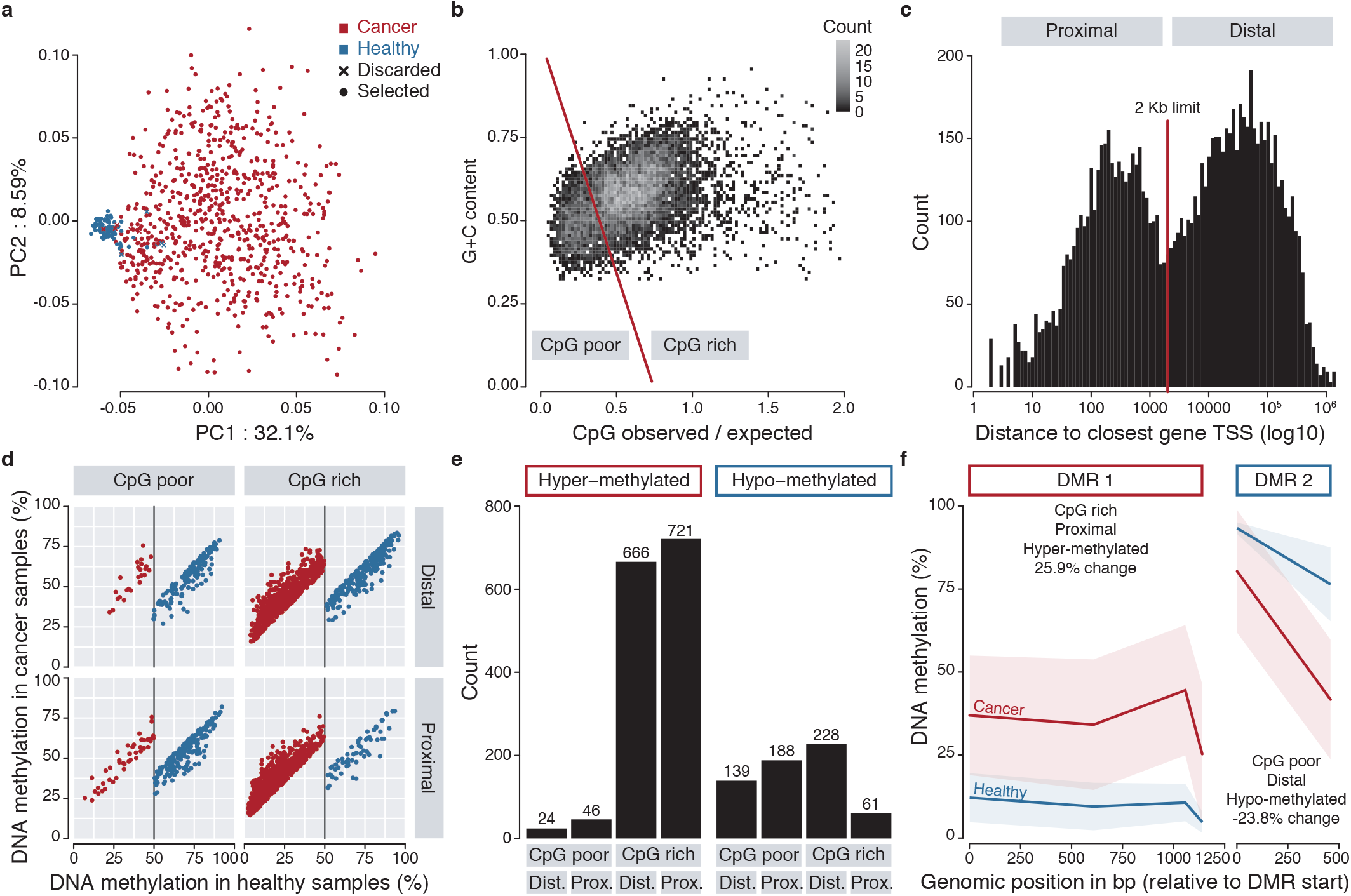
Identification of BRCA DMRs. **a**. Principal component analysis of the BRCA cancer (red) and healthy (blue) samples and visualisation of the samples discarded using a quadratic discriminant analysis (cross). **b**. CpG and G+C content of BRCA DMRs. Two categories, CpG-poor and CpG-rich, were defined according a threshold following y=-1.4(x-0.38)+0.51 **c**. Distance of BRCA DMRs to their closest gene TSS. Two categories, proximal and distal, were defined according to a 2 kilobase (Kb) threshold. **d**. Levels of methylation in hyper-methylated BRCA DMRs (red) or hypo-methylated DMRs (blue) in cancer versus healthy samples in the different categories. Only DMRs with at least 20% methylation change and a starting methylation mean in healthy samples above 50% for DMRs hypo-methylated in cancer and below 50% for hyper-methylated ones are shown. **e**. Number of BRCA hyper- or hypo-methylated DMRs in the different categories. **f**. Example of hyper- and hypo-methylated BRCA DMRs.

### Identification of differentially methylated regions in cancer

We identified DMRs between the cancer and healthy samples for each cancer type by first calling differentially methylated cytosines using limma^17^ and then DMRs using a modified version of DMRcate^18^. Briefly, we only used cytosines not located in exons, which are not expected to contain TF binding sites, and searched for DMRs containing at least two significant CpGs that were consistently hypo- or hyper-methylated.

We further classified DMRs according to their genomic features. Observation of the DMRs’ CpG and G+C contents revealed two distinct categories that we termed CpG-poor and CpG-rich (**Figure 1b**). Observation of the distance of the DMRs to their closest gene transcription start site (TSS) revealed two distinct categories that we termed proximal and distal (**Figure 1c**). These features enable us to minimise the possible biases of the subsequent TF motif analysis due to the preference of some TF to bind specific genomic locations.

We then selected DMRs with a minimum length of 200bp, expected for TF binding sites, at least 20% methylation change and a starting methylation mean in healthy samples above 50% for DMRs hypo-methylated in cancer and below 50% for hyper-methylated ones (**Figure 1d**). Finally, we took advantage of recently available chromatin accessibility ATAC-seq data in TCGA cancer samples^11^ expecting putative TF binding sites to be located in open chromatin ATAC-seq peaks. Therefore, we further selected hypo-methylated DMRs in cancer if they overlapped at least one ATAC-seq peak in the corresponding cancer samples and if hyper-methylated DMRs did not overlap any peak.

The vast majority of DMRs were hyper-methylated and located in CpG-rich regions (**Figure 1e**), which was expected since the TCGA methylation array probes are enriched at gene promoters^19^, usually CpG-rich, and changes in DNA methylation in cancer have previously been described as hyper-methylated in gene promoters. A substantial number of DMRs were also found as hypo-methylated including in CpG-poor regions (**Figure 1e**), which would represent enhancer regulatory regions bound by TFs. Examples of a hyper-methylated CpG-rich proximal DMR i.e. promoter and a hypo-methylated CpG-poor distal DMR i.e. enhancer are shown (**Figure 1f**). The results of these analyses can be visualized interactively for all cancer types on our webserver http://bardet.u-strasbg.fr/cancermethtf/ in the section “Differentially methylated regions”.

### Prediction of transcription factors driving differentially methylated regions in cancer

In order to identify TFs driving DNA methylation changes in cancer, we search for potential TF binding sites in DMRs. TFs bind to DNA through the recognition of short DNA sequences called motifs. We therefore performed an enrichment analysis of known TF motifs using our recently developed approach TFmotifView^20^. We extracted 4928 TF motifs from a manually annotated review^21^, that we could group by similarity into 434 clusters and that represent 1048 distinct TFs. TF motif logos and clusters can be visualised on our webserver http://bardet.u-strasbg.fr/cancermethtf/ in the section “TF motif clusters”. Since hundreds of thousands of TF motifs occurrences can be found over the genome, we computed an enrichment of how many of our DMRs contain at least one occurrence of each motif compared to control regions with similar genomic features or contrasted hypo-methylated DMRs with hyper-methylated DMRs from the same category. We then derived an hypergeometric p-value for each motif enrichment.

We searched for TF motifs in DMRs from all genomic categories and first focused on the hypo-methylated DMRs located in CpG-poor regions distal from genes TSS representing enhancer regions compared to control regions. Across all cancer types, the motifs from the cluster JUN/FOS were highly enriched (although less pronounced for BRCA and PRAD) (**Figure 2a**). Those TFs compose the Activator Protein 1 (AP-1) family that is involved in differentiation, proliferation, apoptosis and well known in tumorigenesis^22^.

**Figure 2:**
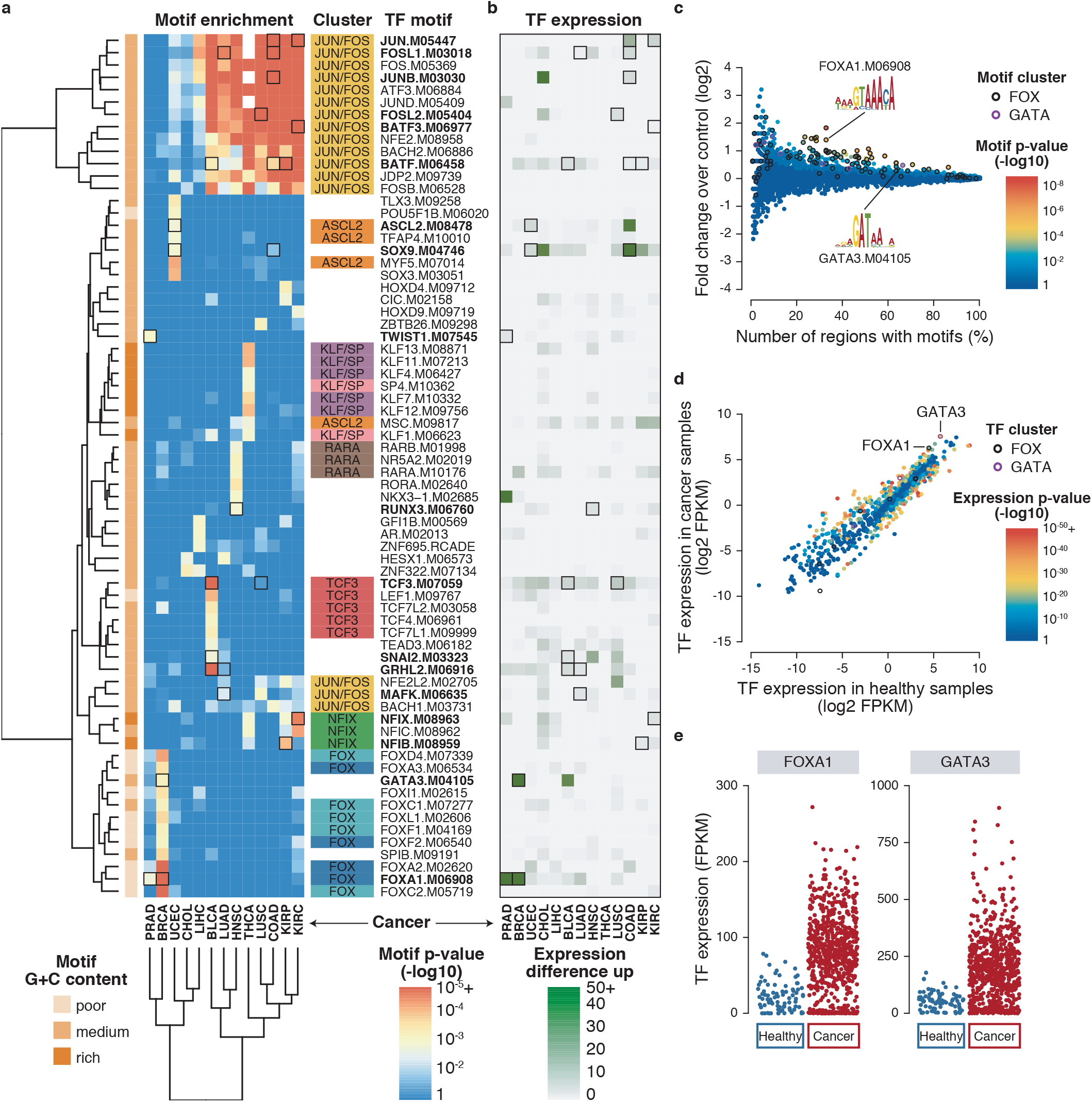
TF motif enrichment in hypo-methylated, CpG-poor, distal DMRs. **a**. Pan cancer motif enrichment. Heatmap of best enriched motifs across all cancer types using a p-value threshold of 10^−3^ and selecting one motif per TF using the p-value sum across all cancers. Motif cluster and CpG content are shown. **b**. Pan cancer TF expression. Heatmap of corresponding TF expression using positive mean FPKM difference between cancer and healthy samples. Motifs in highlighted in bold with black squares have matching motif expression and TF up-regulation. **c**. BRCA motif enrichment. Motif enrichment in BRCA DMRs corresponding to the BRCA column in **a**. Motif p-values (point colour) are computed using an hyper-geometric test using the number of regions that have at least a motif compared to the fold enrichment over control regions. Each point represents one of the 4928 motifs used. FOX and GATA clusters are highlighted (including several FOXA1 or GATA3 motif points). **d**. BRCA TF expression. TF expression enrichment in cancer compared to healthy samples (log2 mean FPKM). Each point represents one of the 1048 TF used coloured according to their differential expression p-value. FOX and GATA clusters are highlighted. **e**. Expression of FOXA1 and GATA3 TFs. Dot plot showing all samples FPKM values for FOXA1 and GATA3 in cancer compared to healthy samples corresponding to the mean value shown in **d**.

Several other TF motifs, sometimes from the same motif cluster and/or TF family, were enriched in specific cancer types, for example the FOX cluster in BRCA (**Figure 2a**). In order to disentangle which specific TF among its motif cluster could drive the cancer DMRs, we integrated matching expression available from TCGA. We hypothesised that hypo-methylated DMRs, methylated in healthy samples and unmethylated in cancer samples, could be regulated in trans by TFs not or lowly expressed in healthy samples and overexpressed in cancer samples therefore binding specifically and driving hypo-methylation in cancer. We then searched for TFs whose expression was upregulated in the cancer samples compared to the healthy samples (**Figure 2b**) and selected TFs that have both their motif enriched and higher expression in cancer (**Figure 2a,b**, black squares). The two most enriched TF motifs with matching upregulation were FOXA1 and GATA3 in BRCA (motif p-value 3.9×10^−6^ and 3.6×10^−4^ respectively; expression upregulation p-value 1.0×10^−21^ and 2.0×10^−28^ respectively). The FOX and GATA motif clusters, containing many putative motifs, represented most of the motifs enriched in BRCA hypo-methylated DMRs located in CpG-poor regions distal from genes TSS (**Figure 2c**). Out of their corresponding TFs, FOXA1 and GATA3 were the most upregulated in BRCA cancer samples (**Figure 2d,e**). Both FOXA1 and GATA3 TFs are known markers in breast cancer^23^.

When searching for motifs enriched in hypo-methylated CpG-poor DMRs either distal or proximal, the following TFs showed both motif enrichment and TF upregulation (**Figure 2a,b and Supplementary Figure 1**): FOXA1, GATA3, RFX5 and TFAP2A in BRCA; TCF3, GRHL2, SNAI2, PATZ1 and BATF in BLCA; JUN, JUNB, FOSL1 and BATF in COAD; RUNX3, FOSL1 and BATF in HNSC; NFIX, BACH1, JUN, BATF and BATF3 in KIRC; NFIB and BATF in KIRP; FOXA1 and FOXA2 in LIHC; HSF1, FOSL1 and BATF in LUAD; NFE2L2, HSF1, GRHL1, GRHL2, TFAP2A, TFAP2C and FOSL2 in LUSC; FOXA1 and TWIST1 in PRAD; ASCL2, SOX9, CEBPB and ESRRA in UCEC. Only very few hyper-methylated DMRs were identified as CpG-poor both distal and proximal and therefore only few TFs showed both motif enrichment and TF downregulation (**Supplementary Figure 1**): KLF5 in BRCA; FOXA2 in CHOL; ARID5B in KIRC; STAT3 in PRAD; RXRG, EGR2 and ZNF263 in UCEC. Many of those TFs have previously been involved in cancer.

When searching for motifs enriched in CpG-rich DMRs, likely to contain different TF motifs due to their distinct sequence content, we contrasted the motif content of hypo-versus hyper-methylated DMRs. Hypo-methylated DMRs were enriched for similar motifs in CpG-rich categories than in CpG-poor categories (**Supplementary Figure 2**). Hyper-methylated CpG-rich DMRs were consistently enriched in G+C-rich low complexity motifs such as EGR1 or KLF in most cancer types (**Figure 3 and Supplementary Figure 2**). EGR1 has been shown to have significant tumor suppressor properties in many types of cancer^24,25^ and different KLF TFs have been involved in a large number of cancers^26^.

**Figure 3:**
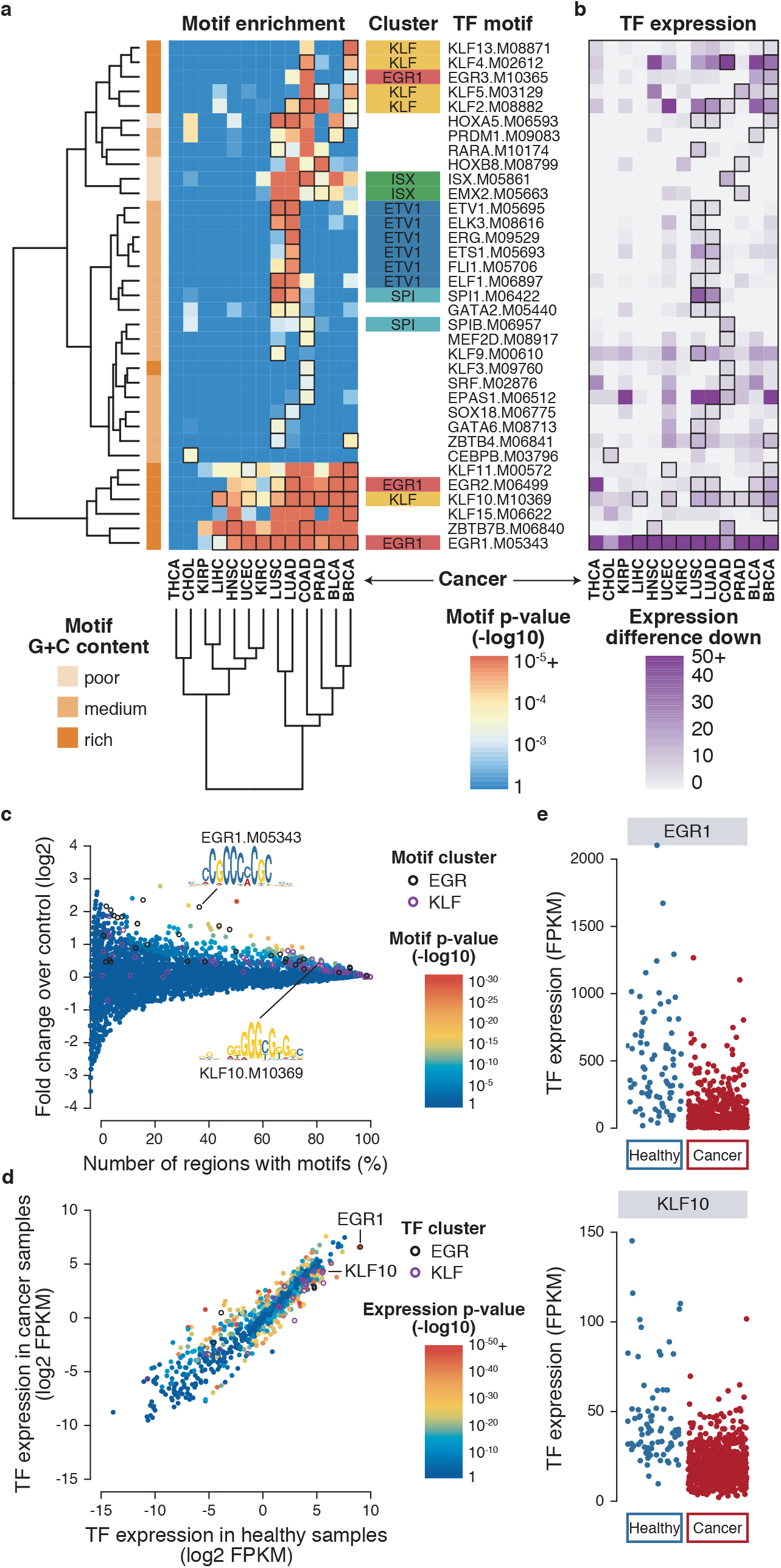
TF motif enrichment in hyper-methylated, CpG-rich, distal DMRs. **a**. Pan cancer motif enrichment. Heatmap of best enriched motifs across all cancer types using a p-value threshold of 10^−3^ and selecting one motif per TF using the p-value sum across all cancers. Only motif with matching expression down-regulation are shown. Motif cluster and CpG content are shown. **b**. Pan cancer TF expression. Heatmap of corresponding TF expression using negative mean FPKM difference between cancer and healthy samples. All motifs represented here have matching expression and TF down-regulation. **C**. BRCA motif enrichment. Motif enrichment in BRCA DMRs corresponding to the BRCA column in **a**. Motif p-values (point colour) are computed using an hyper-geometric test using the number of regions that have at least a motif compared to the fold enrichment over control regions. Each point represents one of the 4928 motifs used. EGR and KLF clusters are highlighted (including several EGR1 or KLF10 motif points). **d**. BRCA TF expression. TF expression enrichment in cancer compared to healthy samples (log2 mean FPKM). Each point represents one of the 1048 TF used coloured according to their differential expression p-value. EGR and KLF clusters are highlighted. **e**. Expression of EGR1 and KLF10 TFs. Dot plot showing all samples FPKM values for EGR1 and KLF10 in cancer compared to healthy samples corresponding to the mean value shown in **d**.

The results of all motif analyses for DMRs in all categories can be visualized interactively for all cancer types on our webserver http://bardet.u-strasbg.fr/cancermethtf/ in the section “TF motif enrichment & expression”.

### Identification of differentially methylated regions in breast cancer cells

We next set out to validate experimentally if the TFs FOXA1 and GATA3 were able to drive changes in DNA methylation in breast cancer cell lines. We used the HCC1954 cell line derived from a primary breast tumor and the hTERT-HME1 cell line as normal mammary epithelial cells. HCC1954 cells show high expression of FOXA1 and GATA3 at both mRNA and protein levels compare to hTERT-HME1 normal cells (**Supplementary Figure 3a,b**) and therefore recapitulate well their expression observed in the TCGA BRCA samples. We performed whole genome bisulfite sequencing (WGBS) on those cell lines and searched for DMRs using DSS^27^ (**Supplementary Table 1 and 2**). We identified 145,826 hypo-methylated DMRs and 121,090 hyper-methylated DMRs in HCC1954 cancer cells compared to hTERT-HME1 normal cells, of which 150 and 1,195 overlapped hypo- and hyper-methylated TCGA BRCA DMRs respectively (**Figure 4a,b**). Additionally, FOXA1 and GATA3 motifs were among the top motifs enriched in all hypo-methylated DMRs compared to hyper-methylated DMRs in all categories (e.g. 1.25 fold enrichment and p-value 0 for FOXA1 and 1.18 fold enrichment and p-value 0 for GATA3 in CpG-poor distal DMRs).

**Figure 4:**
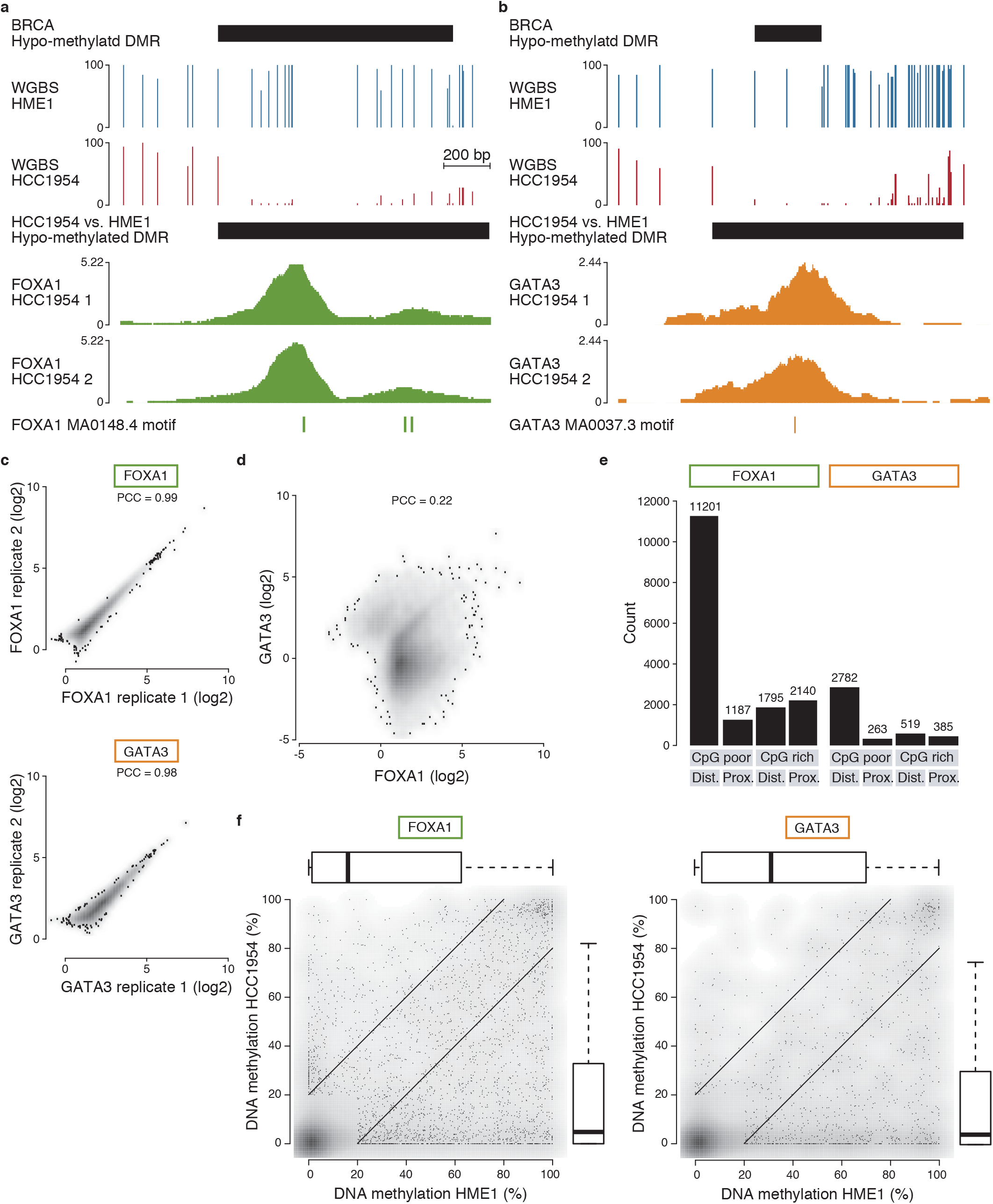
FOXA1 and GATA3 bind hypomethylated regions in HCC1954 breast cancer cells. **a**. Genome browser view (chr14:75521286-75521809) of an hypo-methylated DMR in HCC1954 breast cancer cells compared to hTERT-HME1 normal cells matching a TCGA BRCA hypo-methylated DMR, FOXA1 ChIP-seq signal in HCC1954 cells in two replicates and location of FOXA1 motifs. **b**. Genome browser view (chr2:27209983-27210462) as in **a**. matching a GATA3 binding sites and motif. **c**. Correlation of ChIP-seq signal and Pearson Correlation Coefficient (PCC) of FOXA1 or GATA3 replicates at FOXA1 and GATA3 peaks respectively (FOXA1 n=16323; GATA3 n=3949). **d**. Correlation of ChIP-seq signal and Pearson Correlation Coefficient (PCC) of FOXA1 and GATA3 samples at merged FOXA1 and GATA3 peak regions (n=18270). **e**. Number of FOXA1 and GATA3 binding peaks in the different genomic categories: CpG-poor distal from gene TSS, CpG-poor proximal, CpG-rich distal or CpG-rich proximal. **f**. DNA methylation levels in HCC1954 and hTERT-HME1 in 200bp windows around FOXA1 or GATA3 HCC1954 peak summits that contain at least 2 CpGs and overlapping a matching FOXA1 or GATA3 motif (FOXA1 n=4709; GATA3 n=1671).

### Identification of FOXA1 and GATA3 binding sites in breast cancer cells

We then looked for FOXA1 and GATA3 binding sites in HCC1954 breast cancer cells. We performed chromatin immunoprecipitation sequencing (ChIP-seq) using FOXA1 and GATA3 antibodies in two biological replicates each (**Supplementary Figure 3b and Supplementary Table 3**). After peak calling using peakzilla^28^ and filtering out false positive peaks due to genomic amplification in the HCC1954 breast cancer cells, we obtained 13,753 and 14,257 peaks for FOXA1 replicates samples and 2,095 and 3,751 peaks for GATA3 replicates samples (**Figure 4a,b**). Since the ChIP signal correlated well between replicates (**Figure 4c**), we merged peak regions for further analyses yielding 16,323 peak regions for FOXA1 and 3,949 peak regions for GATA3. Although some binding sites were shared between FOXA1 and GATA3, the majority were distinct (**Figure 4d**). Further, 53% of FOXA1 peak regions and 69% of GATA3 peak regions contained the FOXA1 and GATA3 motifs respectively, which is a usual fraction found in TF ChIP-seq peaks. We next categorized the peak regions according to their genomic features and found that the majority of FOXA1 and GATA3 binding sites were located in CpG-poor regions distal from gene TSS, as expected for TFs to bind distal regulatory regions (**Figure 4e and Supplementary Figure 3c,d**).

Finally, we investigated the methylation patterns at FOXA1 and GATA3 binding sites. We could show that they were located in regions with low DNA methylation levels in cancer HCC1954 cells (**Figure 4f**), expected for TF binding sites in active regulatory regions. We found that 20% and 24% of FOXA1 and GATA3 peak regions respectively overlapped DMRs hypo-methylated in HCC1954 cancer cells compared to hTERT-HME1 normal cells (**Figure 4a,b,f**) and 66 and 35 of FOXA1 and GATA3 peak regions respectively overlapped TCGA BRCA hypo-methylated DMRs (**Figure 4a,b**). They represent putative regions where DNA hypo-methylation could be mediated by FOXA1 or GATA3 binding specifically in cancer cells where they are overexpressed.

### FOXA1 and GATA3 removal in breast cancer cells lead to hyper-methylated DMRs

We next sought to determine if FOXA1 and GATA3 could drive hypo-methylation in HCC1954 breast cancer cells. To test this, we used CRISPR/Cas9 to knockout (KO) FOXA1 or GATA3 in HCC1954 cancer cells (**Figure 5a,b**). We performed WGBS in two independent FOXA1 and GATA3 KO clones (**Supplementary Table 1**) and could observe a gain of DNA methylation in FOXA1 or GATA3 KO cells compared to wildtype (WT) HCC1954 cells at FOXA1 or GATA3 binding sites, respectively (**Figure 5c,d,e,f and Supplementary Figure 4**). We further searched for DMRs independently of FOXA1 or GATA3 binding sites using DSS^27^ (**Supplementary Table 2**) and identified 5969 hyper- and 2578 hypo-methylated DMRs in FOXA1 KO HCC1954 cells compared to WT cells and 14529 hyper-and 1449 hypo-methylated DMRs in GATA3 KO HCC1954 cells. Of those, 84 FOXA1 KO hyper-methylated DMRs overlapped FOXA1 binding peak regions (**Figure 5e**) and 30 GATA3 KO hyper-methylated DMRs overlapped GATA3 binding peak regions (**Figure 5f**), which we expect to result from a direct consequence of FOXA1 or GATA3 removal. We indeed find that FOXA1 or GATA3 binding sites are significantly enriched in FOXA1 or GATA3 KO hyper-over hypo-methylated DMRs (15 fold enrichment with hypergeometric p-value < 10^−14^ for FOXA1 and 2.3 fold with p-value < 10^−7^ for GATA3). This shows that FOXA1 and GATA3 binding do maintain hypo-methylated regions in cancer compared to normal cells.

**Figure 5:**
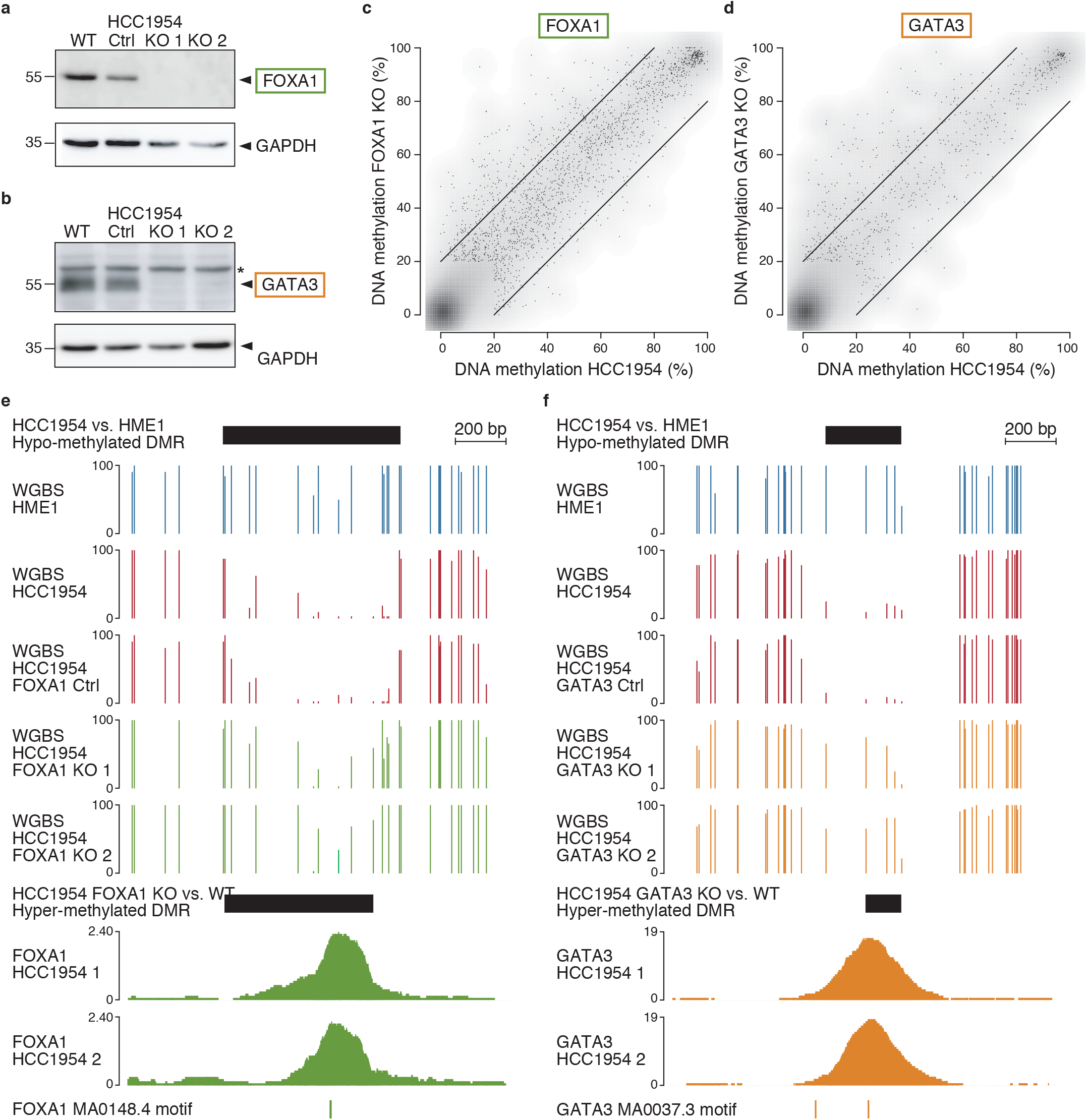
Gain of DNA methylation upon FOXA1 or GATA3 removal in HCC1954 breast cancer cells. **a**. Western blot analysis of FOXA1 protein levels in parental HCC1954 cells (WT), a control clone (ctrl) and two FOXA1 KO clones (KO1 and KO2). GAPDH was used as an internal control for equal loading. **b**. Western blot analysis of GATA3 protein levels in parental HCC1954 cells (WT), a control clone (ctrl) and two GATA3 KO clones (KO1 and KO2). GAPDH was used as an internal control for equal loading. Star indicates nonspecific bands. **c**. DNA methylation levels in HCC1954 FOXA1 KO cells compared to HCC1954 cells (mean across samples) in 200bp windows around FOXA1 peak summits that contain at least 2 CpGs and overlapping a matching FOXA1 motif (n=4473). **d**. DNA methylation levels in HCC1954 GATA3 KO cells compared to HCC1954 cells as in **c**. (n=1598). **e**. Genome browser view (chr19:16093263-16093982) of an hyper-methylated DMR in HCC1954 FOXA1 KO cells compared to HCC1954 cells matching hypo-methylated DMR in HCC1954 vs HME1 and FOXA1 binding sites and motif. **f**. Genome browser view (chr8:42298240-42298413) as in **e**. of an hyper-methylated DMR in HCC1954 GATA3 KO cells compared to HCC1954 cells.

## DISCUSSION

In this study, we took advantage of the massive amount of primary DNA methylation datasets from TCGA and developed a computational approach to predict TFs driving aberrant DNA methylation in cancer. Due to the limited number of CpG probes present in the HM450 array covering 1.7% of the human genome and their bias toward gene promoters^19^, we identified a majority of hyper-methylated DMRs located in CpG-rich regions (**Figure 1e**). However, thanks to the CpG probes designed in putative enhancer regions, we could identify a substantial number of hypo-methylated DMRs (**Figure 1e**).

To predict which TFs could drive those DMRs, we performed a comprehensive TF motif analysis. Since several TFs recognize similar motifs, we used TF gene expression to predict which specific TF could act in trans to mediate DMRs: we searched for downregulated TFs whose motifs were identified in hyper-methylated DMRs and for upregulated TFs in hypo-methylated DMRs. Based on this strategy we did not consider some highly enriched TFs that did not display matching expression such as ZBTB14 in CpG-rich hyper-methylated DMRs in all cancers (except CHOL and THCA), confirming findings from a previous pan-cancer analysis^29^. However, we cannot exclude that TFs expressed at the same level in cancer and normal samples could also impact DNA methylation since this could be due to the change in expression of a partner TF required for the first TF to bind. Additionally, TF motifs enriched in DMRs could arise from TF regulation in cis, where mutations in motifs could affect TF binding leading to DMRs, which we did not investigate in this study.

The results in CpG-rich hyper-methylated DMRs consistently identified CpG-rich low complexity motifs such as EGR1 or KLF motifs in most cancer types (**Figure 3 and Supplementary Figure 2**). However, experimental validations for those TFs, investigating the impact of their loss on DNA methylation, remains challenging since they bind to CpG-rich gene promoters bound by many other TFs that might still maintain the regions unmethylated in their absence. Nevertheless, we do speculate that loss of TF binding does drives hyper-methylation in cancer.

The results in CpG-poor hypo-methylated DMRs predicted TF members of the AP-1 family (motif cluster JUN/FOS) in most cancers (**Figure 2a**), those TFs regulate several important cellular functions such as proliferation, differentiation and apoptosis^30^, and their role in tumorigenesis is well established^22^. In line with this, others studies have previously highlighted an enrichment of the AP-1 motif in hypo-methylated DMRs in colorectal cancer^3,31^.

We also identified other TFs that were more specific to each cancer type **(Figure 2 & Supplementary Figure 1**). Importantly, none of those predicted TFs have prominent CpG in their motifs (besides GRHL1 and CEBPB) making them good candidates to be insensitive to DNA methylation and therefore regulators of DNA methylation patterns^10^. Although many of those TFs are known to be involved in cancer, none have been shown to regulate DNA methylation patterns.

Previous studies have correlated DNA methylation and gene expression changes to identify enhancers and their targets genes^32–34^. ELMER further used 145 TF motifs to infer TF regulators of hypo-methylated TCGA enhancers^33^. Our approach recapitulated several of their predictions such as the AP-1 family TFs (JUN/FOS) in several cancer types, FOXA1 and GATA3 in BRCA, NFE2L2 in LUSC, FOXA2, SOX17, and LEF1 in UCEC, and CEBPB, SPI1 and IRF7 in KIRK **(Supplementary Figures 1 and 2)**.

Based on our computational predictions, which correlate cancer DMRs to TF motif enrichment and differential expression, we chose to validate experimentally if binding of two candidate TFs FOXA1 and GATA3, were indeed upstream of DNA methylation changes. Previous studies also observed a local DNA hypo-methylation at FOXA1 and GATA3 binding sites^35–37^. Both TFs have been described as pioneer TFs as they can bind and open closed chromatin^38^. Moreover, since they do not contain CpG in their motifs, they are less likely to be repressed by DNA methylation^10^, which make them good candidate to drive hypo-methylation in breast cancer.

Due to their pioneer function, they were shown to be involved in hormone-driven cancers by facilitating the access of nuclear receptors to their DNA response elements^38–41^. In breast cancer, FOXA1 and GATA3 are functionally linked with estrogen receptor alpha (ERα), and high level of expression of all three strongly correlates with the luminal subtype of breast tumors also referred as ER positive tumors (ER+)^42,43^. Importantly FOXA1 and GATA3 binding and pioneer function are largely independent of estrogen signaling suggesting an involvement of both TFs in ER negative (ER-) tumors as well^44,45^. We found that BRCA hypo-methylated DMRs called in ER+ or ER-cancer samples compared to healthy samples were both enriched in FOXA1 and GATA3 motifs (motif p-value 0 for both motifs in ER+ DMRs and 9.2×10^−20^ and 1.1×10^−10^ for FOXA1 and GATA3 respectively in ER-DMRs).

Most FOXA1 and GATA3 binding sites in HCC1954 cancer cells were located in low methylated regions (**Figure 4f**). Surprisingly, few were located in fully methylated regions but might be due to the arbitrary definition of the 200bp window around peak summit that might not fit well the specific hypo-methylated region at some loci.

Last we tested the consequence of TF binding on DNA methylation patterns by deleting FOXA1 or GATA3 in HCC1954 cancer cells. Although most changes in DNA methylation did not occur at direct FOXA1 and GATA3 binding sites, which might result from indirect effects, we did observe a gain of DNA methylation at direct FOXA1 or GATA3 binding sites showing that FOXA1 and GATA3 do maintain hypo-methylated regions in cancer cells (**Figure 5**). Although only a limited number of their binding sites gained DNA methylation, it might be explained by binding of other TFs to the same regulatory regions that could maintain the regions unmethylated upon FOXA1 or GATA3 removal. They could be TFs from the same families such as FOXC1, FOXJ2/3, FOXK1/2, FOXM1, FOXN2/3, FOXO1/3/4, FOXP1/4, FOXQ1 or GATA2/6 that are also expressed in HCC1954 cells^4^ or other cooperating TFs.

We did not investigate here the mechanisms by which FOXA1 and GATA3 lead to demethylation and whether this occurs via a passive and/or an active demethylation through the recruitment of Ten-Eleven Translocation (TET) enzymes^46^. Interestingly, FOXA1 was shown to induce TET1 expression through direct binding to its cis-regulatory elements, which in turn led to binding of TET1 to FOXA1 sites mediating local DNA demethylation in prostate cancer cells^47^. However, we do not observe FOXA1 binding sites around the TET1 locus and TET1 is not expressed in HCC1954 breast cancer cells nor in TCGA BRCA samples and other TETs have very low levels of expression.

To summarize, we developed a computational approach to identify TFs driving DNA methylation changes and applied it to TCGA cancer methylation data to predict TFs regulators in 13 different cancer types. This approach could be applied to a wide range of other DNA methylation datasets to infer TF regulators. We validated two TFs, FOXA1 and GATA3 in breast cancer cells, and found that their binding indeed mediate focal hypo-methylation. Altogether this demonstrates the crucial role of TFs in shaping the DNA methylation patterns of a genome and how their deregulation leads to aberrant DNA methylation changes in cancer.

## METHODS

### TCGA methylation data

Raw TCGA Illumina Infinium® HumanMethylation450 BeadChip data was downloaded from the Genomic Data Commons repository (https://gdc.cancer.gov/). Loading of methylation was adapted from the ChAMP pipeline, using the latest HM450 hg38 annotation, removing duplicated CpGs and non-mapping probes. The data was subsequently normalized using noob as implemented in minfi. Principal component analysis (PCA) was then performed on the normalized methylation value of the first 5000 most variable positions, for each cancer. We retained all components whose explained variance is larger than 10% of that of the first component. Quadratic discriminant analysis was then trained on these components, aiming to separate the samples in two classes (cancer and healthy). Samples which were misclassified were discarded.

### Identification of DMRs in TCGA data

Differentially methylated cytosines were called using the R package limma package^17^ at an FDR threshold of 0.05 and excluded cytosines located in exons (using the ENSEMBL annotation for Homo sapiens version GRCh38.87). DMRs were then called using a modified version of DMRcate^18^. The original implementation of DMRcate smoothes a t^2^ statistic and computes p-values using a χ^2^ distribution. This design implies that DMR detection is not sensitive to the sign of methylation change and DMRs could contain a mixture of hypo- or hyper-methylated cytosines. We therefore modified the DMRcate approach to smoothe a t statistic, compute p-values using a normal distribution and used a p-value threshold of 0.001 and parameter λ = 1000. We then selected DMRs or sub-regions of DMRs containing at least 2 consecutive CpGs that were constantly hypo- or hyper-methylated. We then selected DMRs with a minimum length of 200bp, at least 20% methylation change, a starting methylation mean in healthy samples above 50% for DMRs hypo-methylated in cancer and below 50% for hyper-methylated ones and overlapping a corresponding cancer ATAC-seq peak^11^ for hypo-methylated DMRs or not for hyper-methylated ones. We further defined DMRs as CpG-poor or -rich if they were located below or above the line defined by the equation y=-1.4(x-0.38)+0.51 when we compared the ratio of observed versus expected CpG against the G+C content of each DMR. We defined DMRs as proximal or distal if their distance to the closest gene transcriptional start sites was below or above 2000 bp.

### TF motif enrichment

TF motif enrichment was computed and visualised using the TFmotifView approach^20^. Control regions were generated to have the same size than the DMRs, be located in the same genomic context (proximal/distal, CpG-poor/rich, promoter/intron/intergenic), in regions mappable by 50bp reads (non-repetitive), not on chromosome Y and overlapped the same number of HM450 probes. TF motifs were extracted from a manually annotated review^21^, leading to 4928 motifs grouped into 434 clusters (using TOMTOM^48^ and the hclust function in R to define clusters with a threshold of 0.05) and representing 1048 distinct TFs. Using the motif probability matrices we computed for each motif its information content and mean G+C content Let f_ij_ be the frequency of letter i at position j. Then we define *local. IC*_*j*_ = 2 + ∑_*i*_ *f*_*ij*_*log*2(*f*_*ij*_), the information content *IC* = ∑_*j*_ *local. IC*_*j*_, *GC. freq*_*j*_ = *f*_*Gj*_ + *f*_*Cj*_ and mean.GC is the mean of *GC. freq*_*j*_ weighted by its local information content. Motif G+C content was defined as rich (above 0.75), poor (below 0.25) or medium (in between). The hg38 genome was scanned for motif occurrences using mast^49^ with a p-value threshold of 2^-IC^. Motif enrichments were then computed for each DMR category and methylation status by counting the number of DMRs and controls that contain at least one occurrence of a given motif. A pseudocount of 1 was added to all counts. The motif enrichment was defined as the percent of DMRs containing a given motif divided by the percent in control regions. A one-sided hypergeometric p-value was then computed, to test for enrichment significance. Motif enrichments were also computed by comparing, for each category, hypo-methylated versus hyper-methylated DMRs. In that case, the hypergeometric p-value was two-sided to test for depletion as well. Enriched motifs were defined using a p-value threshold of 0.001.

### TCGA expression data

Processed TCGA RNA-seq data was downloaded from the Genomic Data Commons repository (https://gdc.cancer.gov/). FPKMs were averaged for each gene across healthy and cancer samples respectively. Differential expression analysis was performed using DEseq2^50^.

### Webserver for results visualization

The TCGA DMR, TF motif and expression analyses can be visualized on our webserver http://bardet.u-strasbg.fr/cancermethtf/. It was implemented in R using the shiny package (https://shiny.rstudio.com/). It was deployed using the open-source Shiny Server, was containerized using Docker (https://www.docker.com/) and uses Traefik as load-balancer (https://docs.traefik.io/). R shiny servers are optimized for Safari or Microsoft Edge web browsers.

### Cell culture

The cell lines used were obtained from the American Type Culture Collection. The normal mammary epithelial cells, immortalized with hTERT, hTERT-HME1 (ATCC CRL-4010) were cultured in Mammary Epithelial Cell Basal Medium (MEBM) supplemented accordingly to manufacturers (LONZA). HCC1954 cells (ATCC CRL-2328) were cultured in RPMI-1640 medium supplemented with 10% fetal bovine serum and 1% penicillin streptomycin. Cells were maintained in a humidified incubator equilibrated with 5% CO2 at 37 °C. All cell lines were tested negative for mycoplasma.

### Generation of knockout clones

CRISPR/Cas9 genome editing technology was used for generating knockout (KO) cell lines. To disrupt the FOXA1 and GATA3 genes in HCC1954 cells, guide RNAs (gRNAs) targeting the exon 1 of each gene listed in **Supplemental Table 4** were designed by using the Benchling’s CRISPR tool available online (https://benchling.com). gRNAs were synthetized and cloned into the PX459-Puro v2.0 vector (Addgene, # 62988). HCC1954 cells were seeded into six-well plates to achieve 60% confluency before transfection. PX459-gRNA vector was transfected using FuGENE 6 in a 3:1 ratio (µL FuGENE 6: µg DNA) following manufacturer’s instructions. 24h after transfection, transfected cells were transiently growth-selected in medium containing 2 μg/ml puromycin (Gibco) for 48h to eliminate the un-transfected cells. Cells were individually isolated in 96 well plates. Individual clones were further expanded and the loss of FOXA1 or GATA3 expression was confirmed by immunoblotting. One negative clone for FOXA1 or GATA3 were kept and used as controls (Ctrl) in the study.

### RNA isolation, cDNA synthesis and qPCR

Total RNA was extracted using the Allprep DNA/RNA mini kit (Qiagen, catalog #80204). RNA was reverse transcribed using Maxima first strand cDNA synthesis kit (Thermo Fischer Scientific). qPCR was performed with the KAPA SYBR FAST qPCR kit (KAPA Biosystems) on a StepOnePlus PCR system (Applied Biosystems) using the standard curve method. We used fast PCR conditions as follows: 95 °C for 20 s, 40 cycles (95 °C for 20 s, 60 °C for 30 s), followed by a dissociation curve. The expression of target genes was normalized to the RPL13A gene. qPCR reactions were performed in triplicates with no-RT controls to rule out the presence of contaminating DNA. Primers for q-PCR are listed in **Supplementary Table 4**.

### Western blot analysis

Cells were lysed in Pierce™ RIPA Lysis and Extraction buffer (ThermoFisher, #89990) supplemented with protease inhibitors (Roche). The concentration of isolated proteins was determined using Pierce™ BCA protein assay kit (Thermo Fisher, #23227). Protein extracts were run on a 10% SDS polyacrylamide gel and transferred to a 0.2 µm nitrocellulose membrane. The membrane was blocked in TBS, 0.1% Tween-20 containing 5% non-fat dried milk at room temperature for 1 hour and incubated with primary antibodies (dilution 1:1000) at 4°C overnight. The membrane was washed three times with TBS-T, incubated with an appropriate horseradish peroxidase-conjugated secondary antibody for 1 h at room temperature, and washed three times. The signal was detected by chemiluminescence using the ECL detection reagent (Amersham, GE Healthcare). The following primary antibodies were used: anti-FOXA1 (GeneTex, catalog no. GTX100308 and Active motif, catalog no. 39837) and GATA3 (Assay Biotech, catalog no. B0933).

### WGBS

One hundred nanograms of genomic DNA were fragmented to 350 bp using a Covaris E220 sonicator. DNA was bisulfite converted with the EZ DNA Methylation-Gold kit (Zymo Research) and WGBS libraries were prepared using the Accel-NGS Methyl-Seq DNA Library Kit (Swift Biosciences) according to the manufacturer’s instructions with six or seven PCR cycles for the final amplification. The libraries were purified using Ampure XP beads (Beckman Coulter) and sequenced in paired-end (2 × 100 bp) on an Illumina HiSeq4000 at Integragen SA (Evry, France).

### ChIP-seq

Cells were cross-linked with 1% formaldehyde for 8 minutes and quenched by 125mM glycine for 5 minutes at room temperature with gentle shaking. Cells were quickly rinsed in cold PBS twice then scraped in 5mL cold PBS on ice and collected in a 15mL conical tube. Cells were centrifuged at 4°C at 1250xg for 3 minutes. Cell pellets were rinsed with 5mL cold PBS, centrifuged at 4°C at 1250xg for 3 minutes and snap-frozen in liquid nitrogen. Cell pellets were thawed on ice and resuspended in 1mL lysis buffer 1 (50mM HEPES-KOH pH 7.5 140mM NaCl, 1mM EDTA, 10% glycerol, 0.5% NP40, 0.25% Triton X-100) supplemented with protease inhibitors and incubated at 4°C on a rocker for 10 minutes. Lysates were centrifuged at 1000 rpm at 4°C for 5 minutes. Pellets were resuspended with 1mL lysis buffer 2 (10mM Tris pH 8.0 1mM EDTA 0.5mM EGTA 200mM NaCl) supplemented with protease inhibitors and incubated at 4°C on a rocker for 10 minutes. Lysates were centrifuged at 1000 rpm at 4°C for 5 minutes. Pellets were resuspended in 1mL shearing buffer (0.1% SDS, 1mM EDTA, 10mM Tris HCl pH 8.0) supplemented with protease inhibitors, then centrifuged at 1000 rpm at 4°C for 5 minutes. Pellets were resuspended in 500µL shearing buffer, transferred in a 1mL covaris milliTUBE and sonicated with a Covaris E220 sonicator for 8 minutes with 5% duty, 140 peak incident power and 200 cycles per burst. The sonicated lysates were centrifuged at 16000xg for 15 min at 4°C to pellet cellular debris. Sonicated chromatin in the supernatant was transferred to a new 1.5 ml LoBind Eppendorf tube. Immunoprecipitation and elution were performed using the ChIP-IT High Sensitivity kit (Active Motif #53040) according to the manufacturer’s instructions. The following antibodies were used: anti-FOXA1 (GeneTex, catalog no. GTX100308) and GATA3 (Abcam, catalog no. ab199428). Libraries, quality check and sequencing were realized by the GenomEast platform, a member of the “France Génomique” consortium (ANR-10-INBS-0009).

### Sequencing data processing

WGBS reads were trimmed using trim_galore (version 0.6.4 options -q 20 --stringency 2 -- clip_R2 10 --clip_R1 5) (http://www.bioinformatics.babraham.ac.uk/projects/trim_galore/) and mapped using bismark (version 0.22.1)^51^. Non-converted and duplicated reads were further filtered out using filter_non_conversion --percentage_cutoff 50 --minimum_count 5 and deduplicate_bismark. Methylation levels were extracted using bismark_methylation_extractor. DMRs were called using DSS^27^ using CpGs with at least 5 reads coverage as input and selecting DMRs with at least 20% methylation change. For HCC1954 versus HME1 DMRs, we further selected the ones with a minimum length of 200bp and a starting methylation level in HME1 above 50% for hypo-methylated DMRs and below 50% for hyper-methylated DMRs.

ChIP-seq reads were trimmed using trim_galore (version 0.6.4 options -q 20 --stringency 2), mapped using bowtie2 (version 2.3.0)^52^ and selecting reads with mapping quality >= 10. Peaks were called using Peakzilla^28^. In HCC1954 cancer cells, peaks were further filtered out due to localized genomic amplifications. We selected peaks with an input read density lower than its third quartile (0.2438) or that were 10 fold enriched over the input sample.

### Genomic data analyses

All genomic analyses were performed using custom scripts in UNIX using bedtools^53^ and awk and R for plots. Motif enrichment were performed using the JASPAR motifs FOXA1.MA0148.4 and GATA3.MA0037.3.

## DATA AVAILABILITY

Genome-wide sequencing datasets generated for this study are available at NCBI Gene Expression Omnibus under the accession number GSE167870.

## ACKNOWLEDGEMENTS

The authors thank the Weber lab for helpful discussions and Éléa Héberlé for initiating the experiments. This work was funded by the Systems Biology Cancer Plan grant from the ITMO Cancer AVIESAN (French National Alliance for Life Sciences & Health) [18CB008-00 to A.F.B.] and the initiative of excellence IDEX-Unistra from the French national programme Investment for the future [part of the ANR-10-IDEX-0002-02 to A.F.B.]. D.B. is supported by a doctoral fellowship from the Ligue Nationale Contre le Cancer.

## AUTHOR CONTRIBUTIONS

A.F.B conceived and supervised the project. Y.S. performed the bioinformatics predictions. D.D. designed and performed the experiments. D.B. called the DMRs. A.F.B. developed the web application and analysed the genomic data. M.W. provided critical feedback on the project. D.D. and A.F.B. wrote the manuscript and all authors gave feedback on the manuscript.

