## Supplementary material for "Focal DNA hypo-methylation in cancer is mediated by transcription factors binding"

#### Supplementary Table 1: WGBS sequencing data

| Sample | Number raw reads | Percent mapped reads | Percent duplicates removed | Number reads kept | Percent CpG methylation | Mean CpG coverage |
|---|---|---|---|---|---|---|
| WGBS_HME1_1 | 384,099,764 | 83.6% | 8.5% | 293,133,461 | 63.8% | 16 |
| WGBS_HCC1954_1 | 382,607,416 | 82.0% | 25.0% | 235,014,068 | 63.5% | 12 |
| WGBS_HCC1954-FOXA1WT_1 | 445,782,573 | 81.5% | 31.0% | 250,622,961 | 61.5% | 12 |
| WGBS_HCC1954-FOXA1KO_1 | 419,521,228 | 79.0% | 31.0% | 228,549,121 | 64.0% | 11 |
| WGBS_HCC1954-FOXA1KO_2 | 414,620,779 | 77.5% | 31.6% | 219,883,848 | 64.2% | 10 |
| WGBS_HCC1954-GATA3WT_1 | 431,439,156 | 77.6% | 35.5% | 215,281,513 | 61.1% | 10 |
| WGBS_HCC1954-GATA3KO_1 | 447,986,618 | 77.2% | 32.5% | 233,841,837 | 64.0% | 11 |
| WGBS_HCC1954-GATA3KO_2 | 483,415,210 | 77.2% | 32.2% | 252,737,791 | 65.4% | 12 |

#### Supplementary Table 2: DMRs from WGBS data

| Sample 1 | Sample 2 | Number hypo-methylated DMRs | Number hyper-methylated DMRs |
|---|---|---|---|
| WGBS_HCC1954_1 | WGBS_HME1_1 | 145,826 | 121,090 |
| WGBS_HCC1954-FOXA1KO_1<br>WGBS_HCC1954-FOXA1KO_2 | WGBS_HCC1954_1<br>WGBS_HCC1954-FOXA1WT_1 | 2578 | 5969 |
| WGBS_HCC1954-GATA3KO_1<br>WGBS_HCC1954-GATA3KO_2 | WGBS_HCC1954_1<br>WGBS_HCC1954-GATA3WT_1 | 1449 | 14529 |

#### Supplementary Table 3: ChIP-seq sequencing data

| Sample | Number raw reads | Number mapped reads | Percent reads kept | Number peaks | Number merged peaks |
|---|---|---|---|---|---|
| FOXA1_HCC1954_1 | 53,636,539 | 46,336,051 | 86.4% | 13,753 | 16,323 |
| FOXA1_HCC1954_2 | 57,623,146 | 49,609,912 | 86.1% | 14,257 | |
| GATA3_HCC1954_1 | 55,988,365 | 48,262,991 | 86.2% | 2,095 | 3,949 |
| GATA3_HCC1954_2 | 57,303,298 | 49,282,203 | 86.0% | 3,751 | |
| input_HCC1954_1 | 55,828,314 | 32,817,068 | 58.8% | - | - |

#### Supplementary Table 4: ChIP-seq sequencing data

| Gene | Sense | Sequence | Use |
|---|---|---|---|
| FOXA1 | Forward | AGGGCTGGATGGTTGTATTG | qPCR |
| FOXA1 | Reverse | GTGTCTGCGTAGTAGCTGTTC | qPCR |
| GATA3 | Forward | CCAGACCAGAAACCGAAAAA | qPCR |
| GATA3 | Reverse | CCAGACCAGAAACCGAAAAA | qPCR |
| RPL13A | Forward | CCGAGAAGAACGTGGAGAAG | qPCR (housekeeping gene) |
| RPL13A | Reverse | GGCAACGCATGAGGAATTA | qPCR (housekeeping gene) |
| FOXA1 | Forward | CACCGGTAGTAGCTGTTCCAGTCGC | CRISPR/Cas9 (sgRNA) |
| FOXA1 | Reverse | AAACGCGACTGGAACAGCTACTACC | CRISPR/Cas9 (sgRNA) |
| GATA3 | Forward | CACCGGGACTTGCATCCGAAGCCGG | CRISPR/Cas9 (sgRNA) |
| GATA3 | Reverse | AAACCCGGCTTCGGATGCAAGTCCC | CRISPR/Cas9 (sgRNA) |

### Figure S1

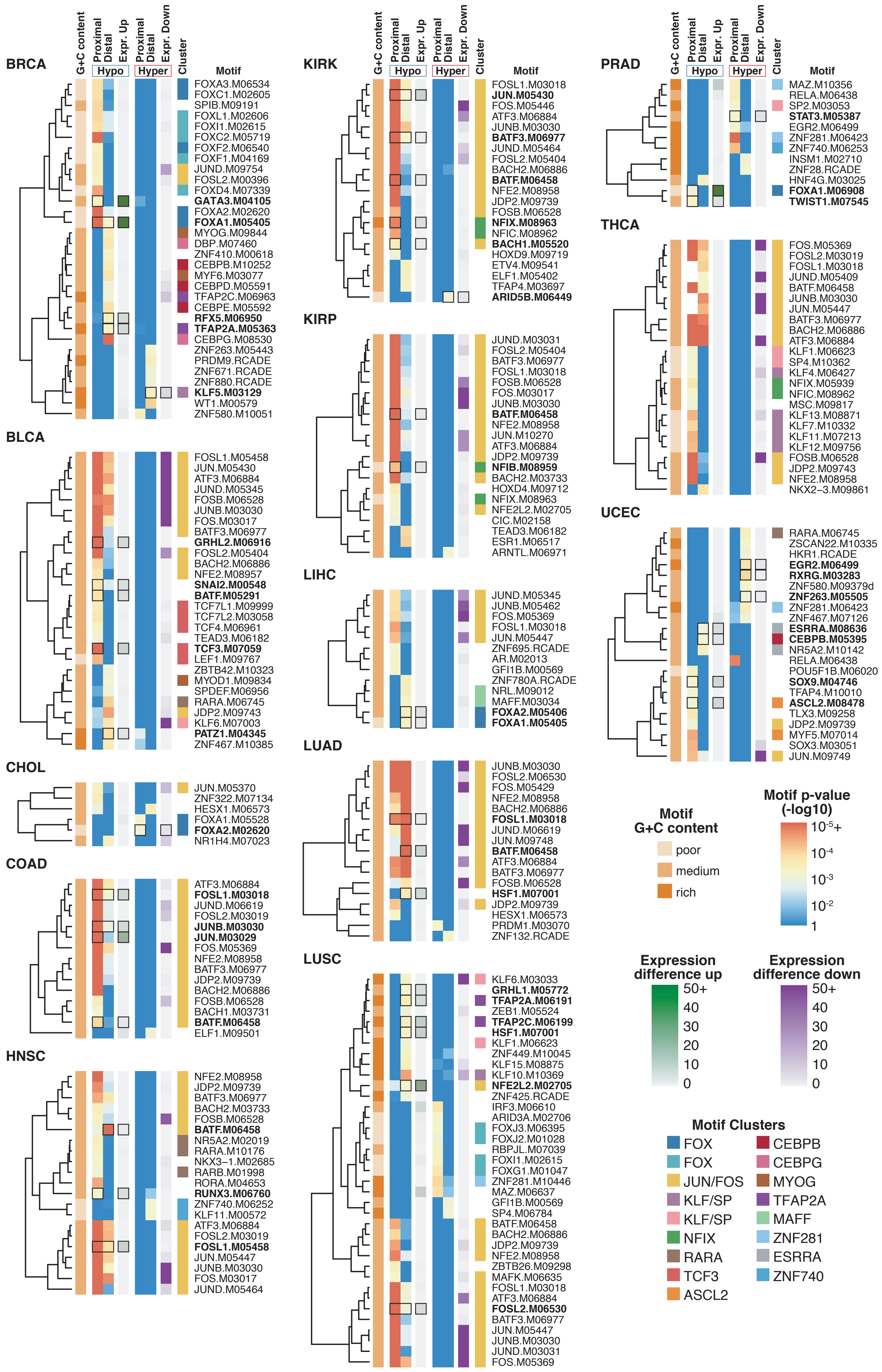

G+C content
Proximal
Distal
Expr. Up
Proximal
Distal
Expr. Down
Cluster
Hypo
Hyper
Motif
BRCA
FOXA3.M06534
FOXC1.M02605
SPIB.M09191
FOXL1.M02606
FOXI1.M02615
FOXC2.M05719
FOXF2.M06540
FOXF1.M04169
JUND.M09754
FOSL2.M00396
FOXD4.M07339
GATA3.M04105
FOXA2.M02620
FOXA1.M05405
MYOG.M09844
DBP.M07460
ZNF410.M00618
CEBPB.M10252
MYF6.M03077
CEBPD.M05591
TFAP2C.M06963
CEBPE.M05592
RFX5.M06950
TFAP2A.M05363
CEBPG.M08530
ZNF263.M05443
PRDM9.RCADE
ZNF671.RCADE
ZNF880.RCADE
KLF5.M03129
WT1.M00579
ZNF580.M10051
BLCA
FOSL1.M05458
JUN.M05430
ATF3.M06884
JUND.M05345
FOSB.M06528
JUNB.M03030
FOS.M03017
BATF3.M06977
GRHL2.M06916
FOSL2.M05404
BACH2.M06886
NFE2.M08957
SNAI2.M00548
BATF.M05291
TCF7L1.M09999
TCF7L2.M03058
TCF4.M06961
TEAD3.M06182
TCF3.M07059
LEF1.M02767
ZBTB42.M10323
MYOD1.M09834
SPDEF.M06956
RARA.M06745
JDP2.M09743
KLF6.M07003
PATZ1.M04345
ZNF467.M10385
CHOL
JUN.M05370
ZNF322.M07134
HESX1.M06573
FOXA1.M05528
FOXA2.M02620
NR1H4.M07023
COAD
ATF3.M06884
FOSL1.M03018
JUND.M06619
FOSL2.M03019
JUNB.M03030
JUN.M03029
FOS.M05369
NFE2.M08958
BATF3.M06977
JDP2.M09739
BACH2.M06886
FOSB.M06528
BACH1.M03731
BATF.M06458
ELF1.M09501
HNSC
NFE2.M08958
JDP2.M09739
BATF3.M06977
BACH2.M03733
FOSB.M06528
BATF.M06458
NR5A2.M02019
RARA.M10176
NKX3−1.M02685
RARB.M01998
RORA.M04653
RUNX3.M06760
ZNF740.M06252
KLF11.M00572
ATF3.M06884
FOSL2.M03019
FOSL1.M05458
JUN.M05447
JUNB.M03030
FOS.M03017
JUND.M05464
KIRK
FOSL1.M03018
JUN.M05430
FOS.M05446
ATF3.M06884
JUNB.M03030
BATF3.M06977
JUND.M05464
FOSL2.M05404
BACH2.M06886
BATF.M06458
NFE2.M08958
JDP2.M09739
FOSB.M06528
NFIX.M08963
NFIC.M08962
BACH1.M05520
HOXD9.M09719
ETV4.M09541
ELF1.M05402
TFAP4.M03697
ARID5B.M06449
KIRP
JUND.M03031
FOSL2.M05404
BATF3.M06977
FOSL1.M03018
FOSB.M06528
FOS.M03017
JUNB.M03030
BATF.M06458
NFE2.M08958
JUN.M10270
ATF3.M06884
JDP2.M09739
NFIB.M08959
BACH2.M03733
HOXD4.M09712
NFIX.M08963
NFE2L2.M02705
CIC.M02158
TEAD3.M06182
ESR1.M06517
ARNTL.M06971
LIHC
JUND.M05345
JUNB.M05462
FOS.M05369
FOSL1.M03018
JUN.M05447
ZNF695.RCADE
AR.M02013
GFI1B.M00569
ZNF780A.RCADE
NRL.M09012
MAFF.M03034
FOXA2.M05406
FOXA1.M05405
LUAD
JUNB.M03030
FOSL2.M06530
FOS.M05429
NFE2.M08958
BACH2.M06886
FOSL1.M03018
JUND.M06619
JUN.M09748
BATF.M06458
ATF3.M06884
BATF3.M06977
FOSB.M06528
HSF1.M07001
JDP2.M09739
HESX1.M06573
PRDM1.M03070
ZNF132.RCADE
LUSC
KLF6.M03033
GRHL1.M05772
TFAP2A.M06191
ZEB1.M05524
TFAP2C.M06199
HSF1.M07001
KLF1.M06623
ZNF449.M10045
KLF15.M08875
KLF10.M10369
NFE2L2.M02705
ZNF425.RCADE
IRF3.M06610
ARID3A.M02706
FOXJ3.M06395
FOXJ2.M01028
RBPJL.M07039
FOXI1.M02615
FOXG1.M01047
ZNF281.M10446
MAZ.M06637
GFI1B.M00569
SP4.M06784
BATF.M06458
BACH2.M06886
JDP2.M09739
NFE2.M08958
ZBTB26.M09298
MAFK.M06635
FOSL1.M03018
ATF3.M06884
FOSL2.M06530
BATF3.M06977
JUN.M05447
JUNB.M03030
JUND.M03031
FOS.M05369
PRAD
MAZ.M10356
RELA.M06438
SP2.M03053
STAT3.M05387
EGR2.M06499
ZNF281.M06423
ZNF740.M06253
INSM1.M02710
ZNF28.RCADE
HNF4G.M03025
FOXA1.M06908
TWIST1.M07545
THCA
FOS.M05369
FOSL2.M03019
FOSL1.M03018
JUND.M05409
BATF.M06458
JUNB.M03030
JUN.M05447
BATF3.M06977
BACH2.M06886
ATF3.M06884
KLF1.M06623
SP4.M10362
KLF4.M06427
NFIX.M05939
NFIC.M08962
MSC.M09817
KLF13.M08871
KLF7.M10332
KLF11.M07213
KLF12.M09756
FOSB.M06528
JDP2.M09743
NFE2.M08958
NKX2−3.M09861
UCEC
RARA.M06745
ZSCAN22.M10335
HKR1.RCADE
EGR2.M06499
RXRG.M03283
ZNF580.M09379d
ZNF263.M05505
ZNF281.M06423
ZNF467.M07126
ESRRA.M08636
CEBPB.M05395
NR5A2.M10142
RELA.M06438
POU5F1B.M06020
SOX9.M04746
TFAP4.M10010
ASCL2.M08478
TLX3.M09258
JDP2.M09739
MYF5.M07014
SOX3.M03051
JUN.M09749
Motif G+C content
poor
medium
rich
Motif p-value (-log10)
10⁻⁵+
10⁻⁴
10⁻³
10⁻²
1
Expression difference up
50+
40
30
20
10
0
Expression difference down
50+
40
30
20
10
0
Motif Clusters
FOX
FOX
JUN/FOS
KLF/SP
KLF/SP
NFIX
RARA
TCF3
ASCL2
CEBPB
CEBPG
MYOG
TFAP2A
MAFF
ZNF281
ESRRA
ZNF740

**Supplementary Figure 1: TF motif enrichment in CpG-poor DMRs**

Pan cancer motif enrichment for all cancer types for all categories of CpG-poor DMRs. Each heatmap shows for one cancer type the best enriched motifs compared to control regions using a p-value threshold of $10^{-3}$ and selecting one motif per TF using the p-value sum across all categories. TF expression heatmap is shown for corresponding TF expression using either positive (up) or negative (down) mean FPKM difference between cancer and healthy samples. Motif cluster and CpG content are shown. Motifs in highlighted in bold with black squares have matching motif expression and TF up- or down-regulation.

**Figure S2**

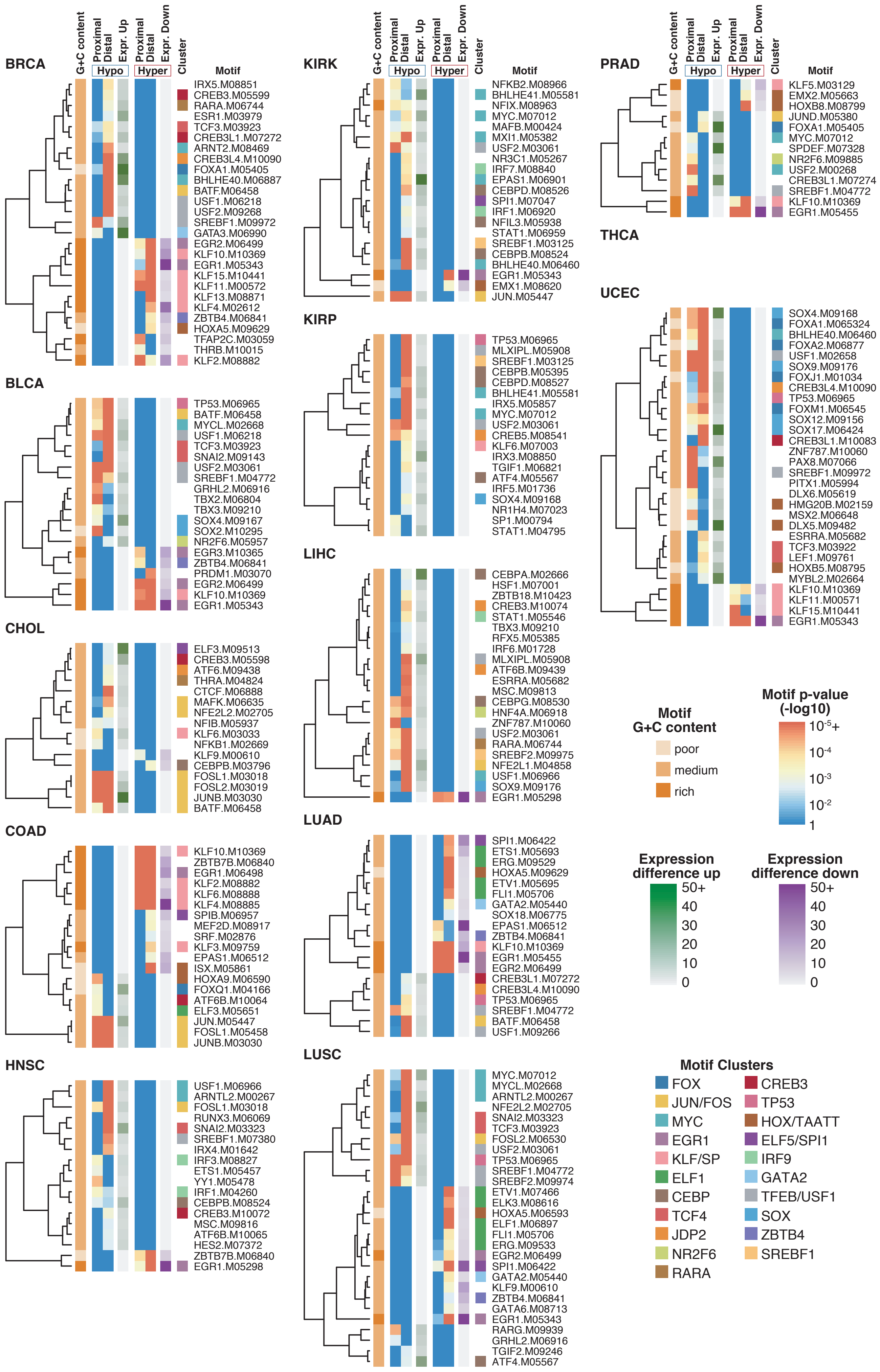
G+C content
Proximal
Distal
Expr. Up
Proximal
Distal
Expr. Down
Hypo
Hyper
Cluster
Motif
BRCA
IRX5.M08851
CREB3.M05599
RARA.M06744
ESR1.M03979
TCF3.M03923
CREB3L1.M07272
ARNT2.M08469
CREB3L4.M10090
FOXA1.M05405
BHLHE40.M06887
BATF.M06458
USF1.M06218
USF2.M09268
SREBF1.M09972
GATA3.M06990
EGR2.M06499
KLF10.M10369
EGR1.M05343
KLF15.M10441
KLF11.M00572
KLF13.M08871
KLF4.M02612
ZBTB4.M06841
HOXA5.M09629
TFAP2C.M03059
THRB.M10015
KLF2.M08882
BLCA
TP53.M06965
BATF.M06458
MYCL.M02668
USF1.M06218
TCF3.M03923
SNAI2.M09143
USF2.M03061
SREBF1.M04772
GRHL2.M06916
TBX2.M06804
TBX3.M09210
SOX4.M09167
SOX2.M10295
NR2F6.M05957
EGR3.M10365
ZBTB4.M06841
PRDM1.M03070
EGR2.M06499
KLF10.M10369
EGR1.M05343
CHOL
ELF3.M09513
CREB3.M05598
ATF6.M09438
THRA.M04824
CTCF.M06888
MAFK.M06635
NFE2L2.M02705
NFIB.M05937
KLF6.M03033
NFKB1.M02669
KLF9.M00610
CEBPB.M03796
FOSL1.M03018
FOSL2.M03019
JUNB.M03030
BATF.M06458
COAD
KLF10.M10369
ZBTB7B.M06840
EGR1.M06498
KLF2.M08882
KLF6.M08888
KLF4.M08885
SPIB.M06957
MEF2D.M08917
SRF.M02876
KLF3.M09759
EPAS1.M06512
ISX.M05861
HOXA9.M06590
FOXQ1.M04166
ATF6B.M10064
ELF3.M05651
JUN.M05447
FOSL1.M05458
JUNB.M03030
HNSC
USF1.M06966
ARNTL2.M00267
FOSL1.M03018
RUNX3.M06069
SNAI2.M03323
SREBF1.M07380
IRX4.M01642
IRF3.M08827
ETS1.M05457
YY1.M05478
IRF1.M04260
CEBPB.M08524
CREB3.M10072
MSC.M09816
ATF6B.M10065
HES2.M07372
ZBTB7B.M06840
EGR1.M05298
KIRK
NFKB2.M08966
BHLHE41.M05581
NFIX.M08963
MYC.M07012
MAFB.M00424
MXI1.M05382
USF2.M03061
NR3C1.M05267
IRF7.M08840
EPAS1.M06901
CEBPD.M08526
SPI1.M07047
IRF1.M06920
NFIL3.M05938
STAT1.M06959
SREBF1.M03125
CEBPB.M08524
BHLHE40.M06460
EGR1.M05343
EMX1.M08620
JUN.M05447
KIRP
TP53.M06965
MLXIPL.M05908
SREBF1.M03125
CEBPB.M05395
CEBPD.M08527
BHLHE41.M05581
IRX5.M05857
MYC.M07012
USF2.M03061
CREB5.M08541
KLF6.M07003
IRX3.M08850
TGIF1.M06821
ATF4.M05567
IRF5.M01736
SOX4.M09168
NR1H4.M07023
SP1.M00794
STAT1.M04795
LIHC
CEBPA.M02666
HSF1.M07001
ZBTB18.M10423
CREB3.M10074
STAT1.M05546
TBX3.M09210
RFX5.M05385
IRF6.M01728
MLXIPL.M05908
ATF6B.M09439
ESRRA.M05682
MSC.M09813
CEBPG.M08530
HNF4A.M06918
ZNF787.M10060
USF2.M03061
RARA.M06744
SREBF2.M09975
NFE2L1.M04858
USF1.M06966
SOX9.M09176
EGR1.M05298
LUAD
SPI1.M06422
ETS1.M05693
ERG.M09529
HOXA5.M09629
ETV1.M05695
FLI1.M05706
GATA2.M05440
SOX18.M06775
EPAS1.M06512
ZBTB4.M06841
KLF10.M10369
EGR1.M05455
EGR2.M06499
CREB3L1.M07272
CREB3L4.M10090
TP53.M06965
SREBF1.M04772
BATF.M06458
USF1.M09266
LUSC
MYC.M07012
MYCL.M02668
ARNTL2.M00267
NFE2L2.M02705
SNAI2.M03323
TCF3.M03923
FOSL2.M06530
USF2.M03061
TP53.M06965
SREBF1.M04772
SREBF2.M09974
ETV1.M07466
ELK3.M08616
HOXA5.M06593
ELF1.M06897
FLI1.M05706
ERG.M09533
EGR2.M06499
SPI1.M06422
GATA2.M05440
KLF9.M00610
ZBTB4.M06841
GATA6.M08713
EGR1.M05343
RARG.M09939
GRHL2.M06916
TGIF2.M09246
ATF4.M05567
PRAD
KLF5.M03129
EMX2.M05663
HOXB8.M08799
JUND.M05380
FOXA1.M05405
MYC.M07012
SPDEF.M07328
NR2F6.M09885
USF2.M00268
CREB3L1.M07274
SREBF1.M04772
KLF10.M10369
EGR1.M05455
THCA
UCEC
SOX4.M09168
FOXA1.M065324
BHLHE40.M06460
FOXA2.M06877
USF1.M02658
SOX9.M09176
FOXJ1.M01034
CREB3L4.M10090
TP53.M06965
FOXM1.M06545
SOX12.M09156
SOX17.M06424
CREB3L1.M10083
ZNF787.M10060
PAX8.M07066
SREBF1.M09972
PITX1.M05994
DLX6.M05619
HMG20B.M02159
MSX2.M06648
DLX5.M09482
ESRRA.M05682
TCF3.M03922
LEF1.M09761
HOXB5.M08795
MYBL2.M02664
KLF10.M10369
KLF11.M00571
KLF15.M10441
EGR1.M05343
Motif
G+C content
poor
medium
rich
Motif p-value
(-log10)
10^-5+
10^-4
10^-3
10^-2
1
Expression
difference up
50+
40
30
20
10
0
Expression
difference down
50+
40
30
20
10
0
Motif Clusters
FOX
JUN/FOS
MYC
EGR1
KLF/SP
ELF1
CEBP
TCF4
JDP2
NR2F6
RARA
CREB3
TP53
HOX/TAATT
ELF5/SPI1
IRF9
GATA2
TFEB/USF1
SOX
ZBTB4
SREBF1

**Supplementary Figure 2: TF motif enrichment in CpG-rich DMRs**

Pan cancer motif enrichment for all cancer types for all categories of CpG-rich DMRs. Each heatmap shows for one cancer type the best enriched motifs in hypo- compared to hyper-methylated regions using a p-value threshold of $10^{-3}$ and selecting one motif per TF using the p-value sum across all categories. TF expression heatmap is shown for corresponding TF expression using either positive (up) or negative (down) mean FPKM difference between cancer and healthy samples. Only motifs that have matching motif expression and TF up- or down-regulation are shown. Motif cluster and CpG content are shown.

### Figure S3

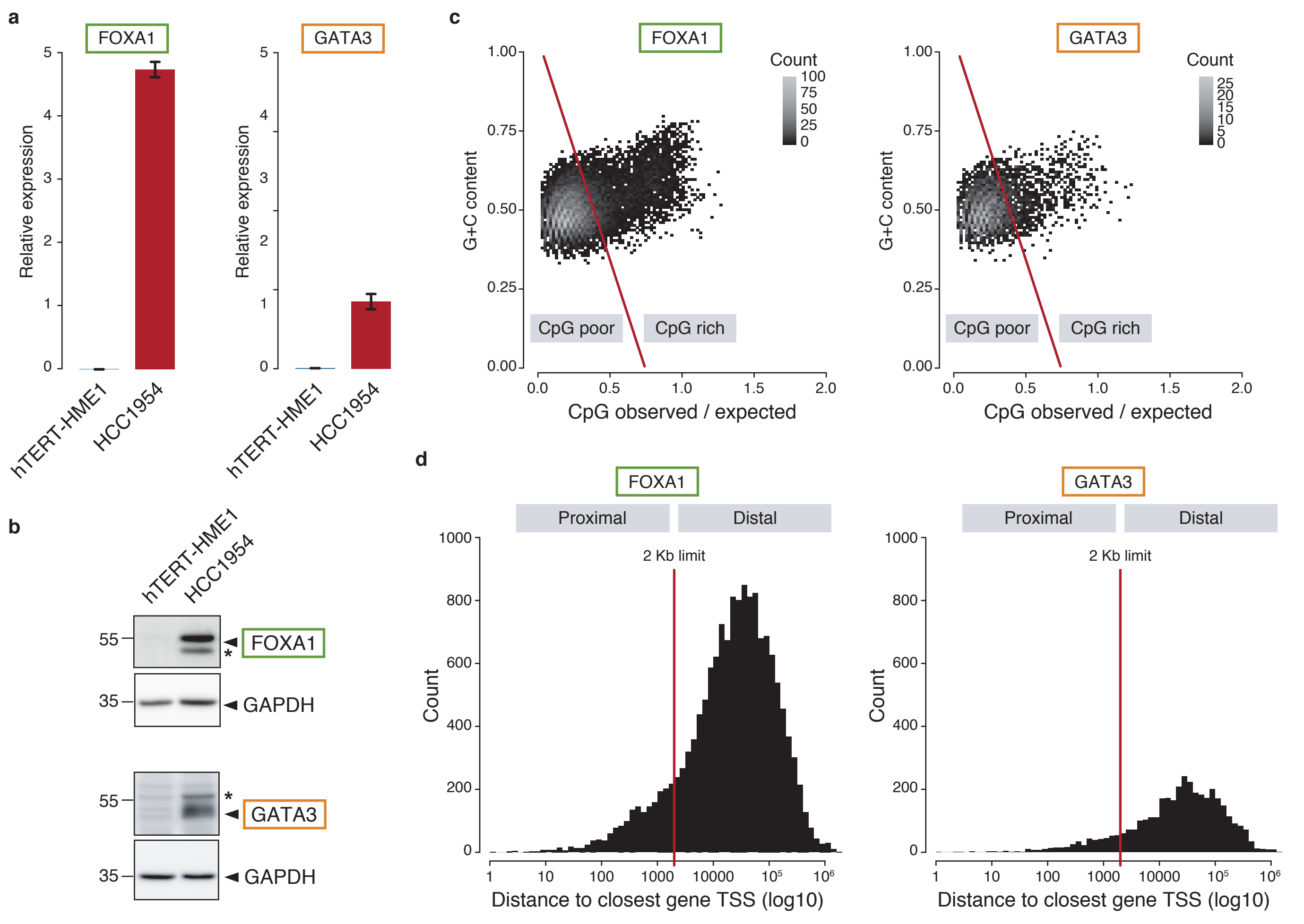

a
FOXA1
GATA3
Relative expression
hTERT-HME1
HCC1954
b
hTERT-HME1
HCC1954
55
35
FOXA1
GAPDH
GATA3
c
FOXA1
GATA3
Count
G+C content
CpG poor
CpG rich
CpG observed / expected
d
Proximal
Distal
2 Kb limit
Count
Distance to closest gene TSS (log10)

**Supplementary Figure 3: FOXA1 and GATA3 expression and binding in HCC1954 breast cancer cells**

**a.** Expression levels of FOXA1 and GATA3 in hTERT-HME1 and HCC1954 cells by RT-qPCR (mean +/- SEM, n=3; relative to RPL13A expression).

**b.** Protein levels of FOXA1 and GATA3 in hTERT-HME1 and HCC1954 cells by western blotting. GAPDH was used as an internal control for equal loading. Stars indicate non-specific bands.

**c.** CpG and G+C content of FOXA1 and GATA3 peaks. Two categories, CpG-poor and CpG-rich, were defined according a threshold following $y=-1.4(x-0.38)+0.51$ (FOXA1 n=16323; GATA3 n=3949).

**d.** Distance of FOXA1 and GATA3 peaks to their closest gene TSS. Two categories, proximal and distal, were defined according to a 2 kilobase (Kb) threshold (FOXA1 n=16323; GATA3 n=3949).

#### Figure S4

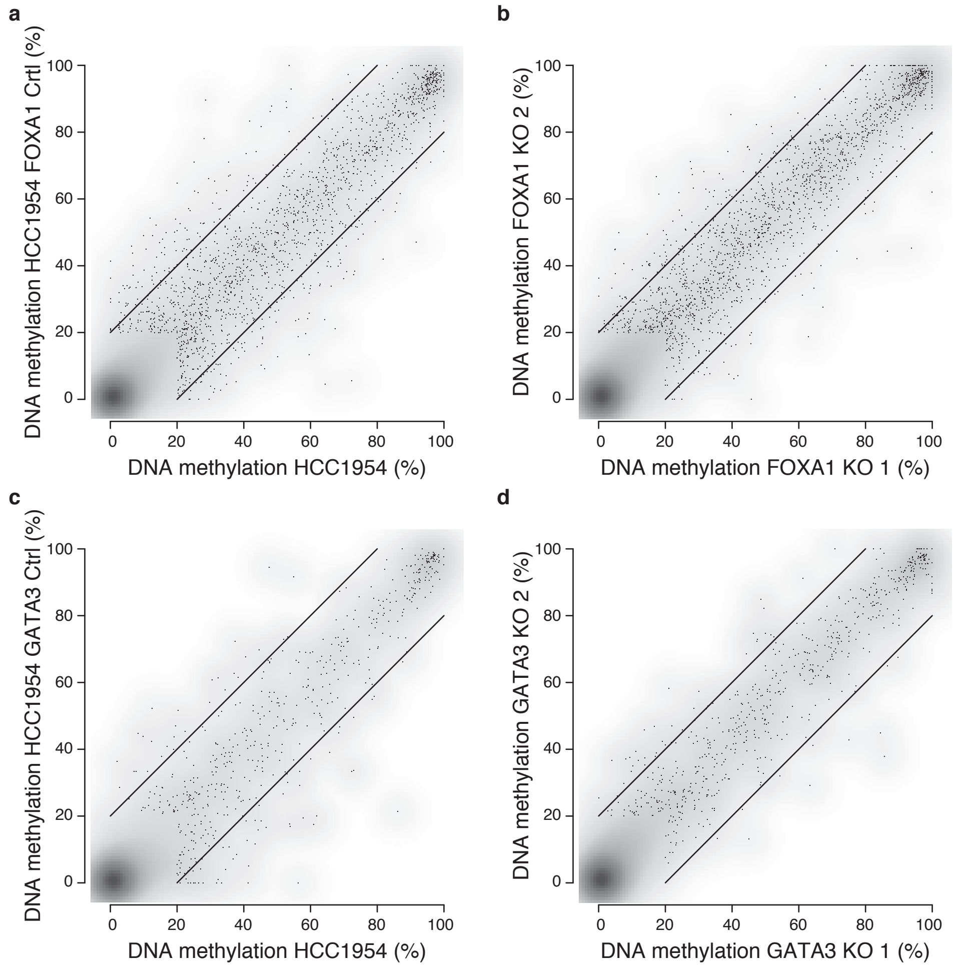
a
b
c
d
DNA methylation HCC1954 FOXA1 Crtl (%)
DNA methylation HCC1954 (%)
DNA methylation FOXA1 KO 2 (%)
DNA methylation FOXA1 KO 1 (%)
DNA methylation HCC1954 GATA3 Ctrl (%)
DNA methylation HCC1954 (%)
DNA methylation GATA3 KO 2 (%)
DNA methylation GATA3 KO 1 (%)
0
20
40
60
80
100

**Supplementary Figure 4: DNA methylation levels in HCC1954 breast cancer cells**

**a.** DNA methylation levels in WT HCC1954 and HCC1954 FOXA1 KO control cells (mean across samples) in 200bp windows around FOXA1 peak summits that contain at least 2 CpGs and overlapping a matching FOXA1 motif (n=4473).

**b.** DNA methylation levels in two replicates of HCC1954 FOXA1 KO cells as in **a**.

**c.** DNA methylation levels in WT HCC1954 and HCC1954 GATA3 KO control cells as in **a**. and overlapping a matching GATA3 motif (n=1598).

**d.** DNA methylation levels in two replicates of HCC1954 GATA3 KO cells as in **c**.
